# A synthetic kinematic index of trunk displacement conveying the overall motor condition in Parkinson’s disease

**DOI:** 10.1101/2020.07.13.199224

**Authors:** Emahnuel Troisi Lopez, Roberta Minino, Pierpaolo Sorrentino, Rosaria Rucco, Anna Carotenuto, Valeria Agosti, Domenico Tafuri, Valentino Manzo, Marianna Liparoti, Giuseppe Sorrentino

## Abstract

**BACKGROUND:** Parkinson’s disease (PD) is characterized by motor impairment, affecting quality of life and increasing fall risk, due to ineffective postural control. To this day, the diagnosis remains based on clinical approach. Similarly, motor evaluation is based on heterogeneous, operator-dependent observational criteria. A synthetic, replicable index to quantify motor impairment is still lacking. In this paper, we build upon the idea that the trunk is crucial in balance control. Hence, we have designed a new measure of postural stability which assesses the trunk displacement in relation to the center of mass, that we named trunk displacement index (TDI).

**METHODS:** Twenty-three PD patients and twenty-three healthy controls underwent clinical (UPDRS-III) and motor examination (3D gait analysis). The TDI was extracted from kinematic measurements using a stereophotogrammetric system. A correlation analysis was performed to assess the relationship of TDI with typical gait parameters, to verify its biomechanical value, and UPDRS-III, to observe its clinical relevance. Finally, its sensitivity was measured, comparing pre- and post-L-DOPA subclinical intake.

**RESULTS:** The TDI showed significant correlations with many gait parameters, including both velocity and stability characteristics of gait, and with the UPDRS-III. Finally, the TDI resulted capable in discriminating between off and on state in PD, whereas typical gait parameters failed two show any difference between those two conditions.

**CONCLUSIONS:** Our results suggest that the TDI may be considered a highly sensitive biomechanical index, reflecting the overall motor condition in PD, and provided of clinical relevance due to the correlation with the clinical evaluation.

## 1 INTRODUCTION

Parkinson’s disease (PD), the second most common progressive neurodegenerative disease, is characterized primarily by degeneration of the dopamine-secreting neurons in the pars compacta of the substantia nigra^1^. While PD occurs with both motor and non-motor symptoms the motor phenotype has been specifically linked to the quality of life^2^. More specifically, gait alterations contribute to balance impairment and, hence, increased risk of falling^3–5^. Oral therapy with L-DOPA is currently the gold-standard treatment for the disease^6^. In order to assess the clinical state and the disease progression, the most widely used rating scale is the Unified Parkinson’s Disease Rating Scale (UPDRS)^7^. The UPDRS part III concerns the evaluation of the motor signs and provides a score representing the motor impairment of PD patients. However, UPDRS-III suffers of several limitations, being its scoring based on a summation of subjective evaluation of heterogeneous motor features. An objective way to analyse movement might be of help to overcome such limitations.

Three-dimensional motion analysis (3D-MA) is a quantitative method to detect movements with high spatial resolution, and is regarded as the gold standard for movement evaluation^8^. 3D-MA is widely used for the assessment of motor skills and since locomotion is importantly affected in PD a great deal of analyses have been specifically devoted to analysis of gait^9–11^. However, most of the studies using 3D-Gait Analysis (3D-GA) in PD focused on lower limbs parameters such as step length, stride and swing time, range of motion (ROM) of joints^12–14^, dismissing other body segments. Consequently, despite extensive efforts retrieving a synthetic and informative parameter able to reflect the overall motor condition has proven elusive.

In this study, we inspected the following line of reasoning, in order to retrieve a synthetic and informative parameter to assess gait stability. From an evolutionistic perspective, locomotion is a complex process, and the lower limbs are part of a highly-structured kinematic chain that includes all body segments. In the evolution of upright posture and bipedal locomotion, the position of the trunk changed, and so did the distribution of its weight on the lower limbs^15,16^. Such changes provoked a marked rise in the complexity of the task of keeping balance^17^ becoming computationally demanding. To this regard, suprasegmental control has been shown to add considerable stability while reducing computational load. In fact, in order to keep balance, the cerebellum control does not operate on each muscle separately, but rather it aims to the control of the centre of mass(COM), specifically integrating information with vestibulospinal information of trunk verticality^17,18^.

Following this evolutionary reasoning, we took into consideration the well-known PD trunk impairment and the relative increased fall risk^19^. Intuitively, a key aspect facilitating the stability of the centre of mass is that the pectoral girdle should not be oscillating too much as compared to the COM, otherwise the subject would be more prone to falls. However, 3D-GA studies that appraise the trunk movement are limited. Recently, Bestaven et al., through 3D-GA in ten PD patients before and after rehabilitation, performed a trunk analysis measuring the horizontal distance between C7 and L3 vertebrae and the vertical distance between acromia. Despite a trend in the latter, both results were not statistically different^20^. In 2010, Roiz et al., through 3D-GA, measured trunk flexion on the sagittal plane in 12 PD patients and 15 healthy controls, with no statistically significant results^21^.

The aim of our study is to find an objective biomechanical index to synthetically convey the effect of the complex motor impairment in PD on stability. We introduced a new index, which quantifies the trunk displacement in relation to the COM. The index values were obtained from 3D-GA data, evaluating the ratio of the displacements of trunk and COM, on three anatomical planes, in 23 early PD patients, during gait. To evaluate the ability of the trunk displacement index (TDI) to effectively synthetize many different motor characteristics, we carried out a correlation analysis between TDI and typical 3D-GA parameters. The same analysis was performed between TDI and UPDRS in order to evaluate the association of TDI with clinical motion condition. Finally, to test the sensitivity of the TDI to detect slight changes in motion features, we compared its value in the PD patient before and after a subclinical (half of the morning dose) L-DOPA intake.

## 2 METHODS

### 2.1 Participants

Twenty-three in-patients (PD group), referred to the Movement Disorders Unit of the Cardarelli Hospital in Naples were recruited. PD diagnosis was defined according to the United Kingdom Parkinson’s Disease Brain Bank criteria^22^. Inclusion criteria were: a) minimum age of 45 years or older; b) Hoehn & Yahr (H&Y) score ≤ 3 while at “off” state; c) disease duration < 10 years; d) antiparkinsonian treatment at a stable dosage. Exclusion criteria were: a) Mini-Mental State Examination (MMSE) < 24^23^; b) Frontal Assessment Battery (FAB) < 12^24^; c) Beck Depression Inventory II (BDI-II) > 13^25^; d) neurological (except PD) or psychiatric disorders; e) assumption of psychoactive drugs; f) any physical or medical conditions causing walking impairment. Twenty-three healthy people matched for age, gender, education has been recruited as control group (HC group). Exclusion criteria were the same of the PD group.

According to the declaration of Helsinki, an informed consent was obtained from all participants. The study was approved by the AORN “A. Cardarelli” Ethic Committee.

### 2.2 Intervention

Participants were asked to walk in the laboratory at self-selected speed, in a straight path. The control group was recorded once. Conversely, the PD group was acquired twice: during the first acquisition the patients were in off state (no L-DOPA intake in the last 14-16 hours) (PDoff group), while in the second acquisition the subjects were recorded 40min after taking a subclinical dose (defined as half of their usual morning intake) of L-DOPA (Melevodopa + Carbidopa) (PDon group). Before each acquisition, the PD subjects were tested through UPDRS-III. For each participant and each condition (off and on in PD group), four gait cycles were acquired and averaged for data analysis.

### 2.3 Motion analysis

#### 2.3.1 Acquisition system

The gait analysis took place in the Motion Analysis Laboratory of the University of Naples Parthenope. The motion capture has been achieved through a stereophotogrammetric system consisting of eight infrared cameras (ProReflex Unit - Qualisys Inc., Gothenburg, Sweden). Fifty-five markers were applied on each participant according to the modified Davis protocol^26^ on anatomical landmarks of feet, lower limbs joints, pelvis, trunk, upper limbs joints and head. Before the trials, all participants were instructed to walk at a normal pace through the measured space (10m). Through 3D-GA, the following typical gait parameters were calculated and corrected for the body mass index (BMI).

#### 2.3.2 Spatiotemporal parameters

Spatio-temporal parameters were divided in two categories, representing velocity and stability characteristics of gait^27^. The velocity parameters included speed (meters/second), stride length (meters), cadence (strides/minute), cycle time (seconds). The stability parameters consisted of stride width (meters), stance time (seconds), swing time (seconds), double limb support time (DLS) (seconds). For each spatio-temporal value (excluding speed) we computed the CV, a derived parameter expression of stability^28^, obtained by dividing the standard deviation by the mean value, then multiplying the result by one hundred.

#### 2.3.3 Kinematic parameters

Furthermore, kinematics parameters of the lower limb joints (ankle (A), knee (K), thigh (T)) were calculated to specifically collect the angular values of each flexion/extension peak, normalized for the 100% of the gait cycle ^29,30^. Then, we calculated the difference between each consecutive peak, and obtained 4 Δ values (Figure 1) for each joint, representative of the angular excursion during the gait cycle (AΔ1-2-3-4, KΔ1-2-3-4, TΔ1-2-3-4).

**Figure 1.**
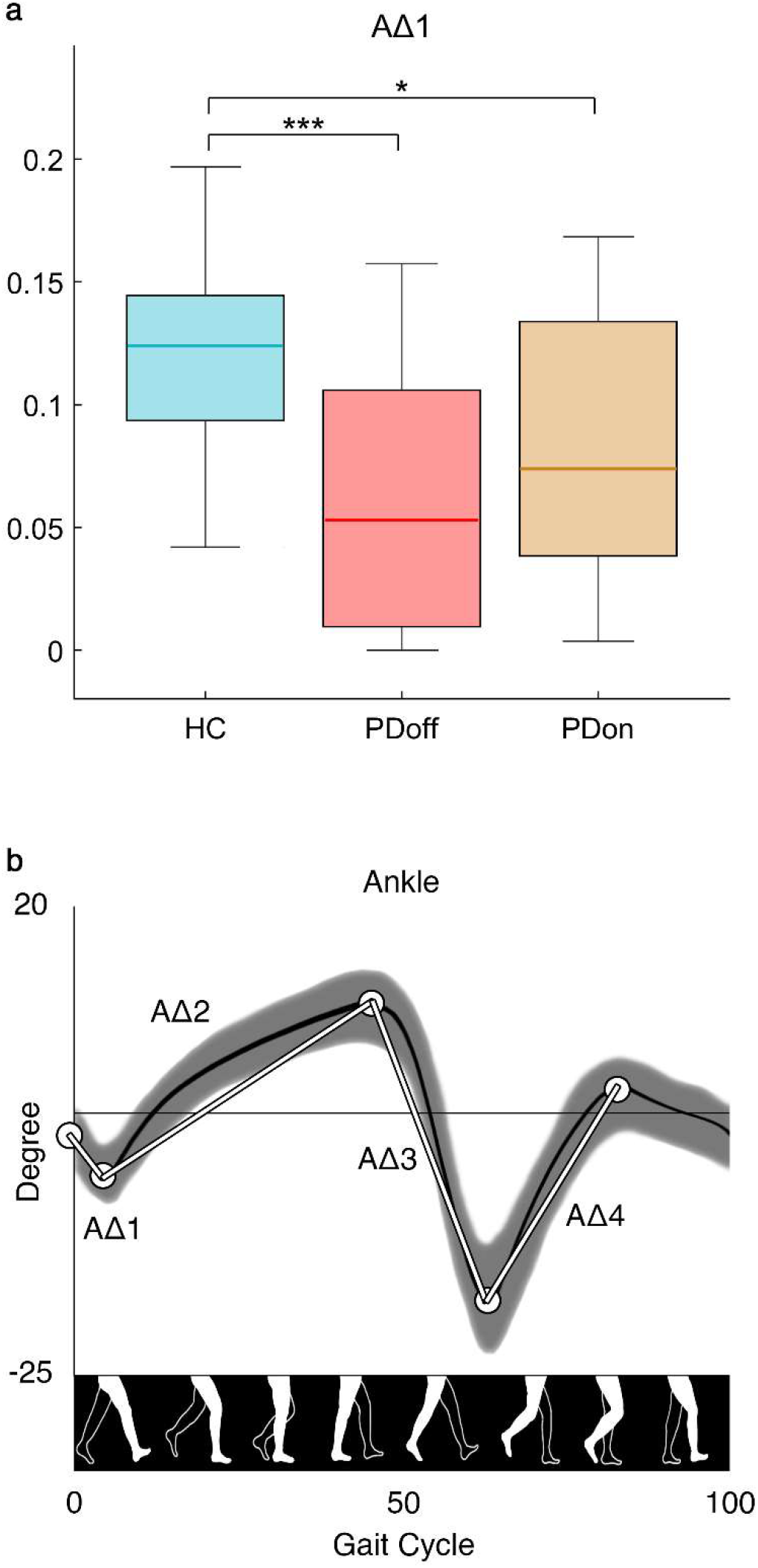
Kinematic analysis of gait. **a**. The box plot of the ankle excursion during the initial phase of gait cycle (AΔ1). The Δ is calculated as the difference between two consecutive peaks. The box plot includes data from 25th to 75th percentiles; the median is represented by the horizontal line inside each box; error lines reach the 10th and 90th percentiles; the outliers, if present, are represented by filled circles falling beyond 10th and 90th percentiles. Significance *p* value: \**p*<0.05, \*\**p*<0.01, \*\*\**p*<0.001. Healthy controls (HC), individual with Parkinson’s disease before L-DOPA intake (PDoff), individual with Parkinson’s disease after L-DOPA intake (PDon). **b**. The traces of the normal kinematic curves. The differences between peaks (Δs) are represented by the white lines.

#### 2.3.4 Trunk Displacement Index

Beyond typical gait parameters, according to our initial hypothesis, we designed a new measurement method to evaluate the trunk displacement. Firstly, we calculated the 3D trajectory of the centre of mass *COMt* of each subject and its mean 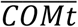 during gait. Then we looked for an anatomical reference point, representative of the trunk position and located it between acromia. As for the COM, we calculated the trunk trajectory *Tt*. Then, we calculated the distances of each point of *COMt* and *Tt* from 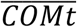, respectively named *COMd* and *Td*, during the whole gait cycle as (Figure 2a):

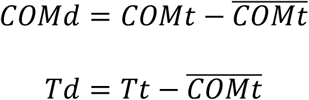

obtaining their respective vectors of distances in all three planes. To enclose the three-dimensional data in a unique value, for each vector of distances we calculated its norm and summed the results separately. Finally, we measured the ratio between the two values to capture the relationship between those two segments, in the form of a dimensionless quantity (from now on called trunk displacement index or TDI).

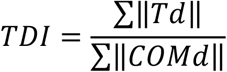

**Figure 2.**
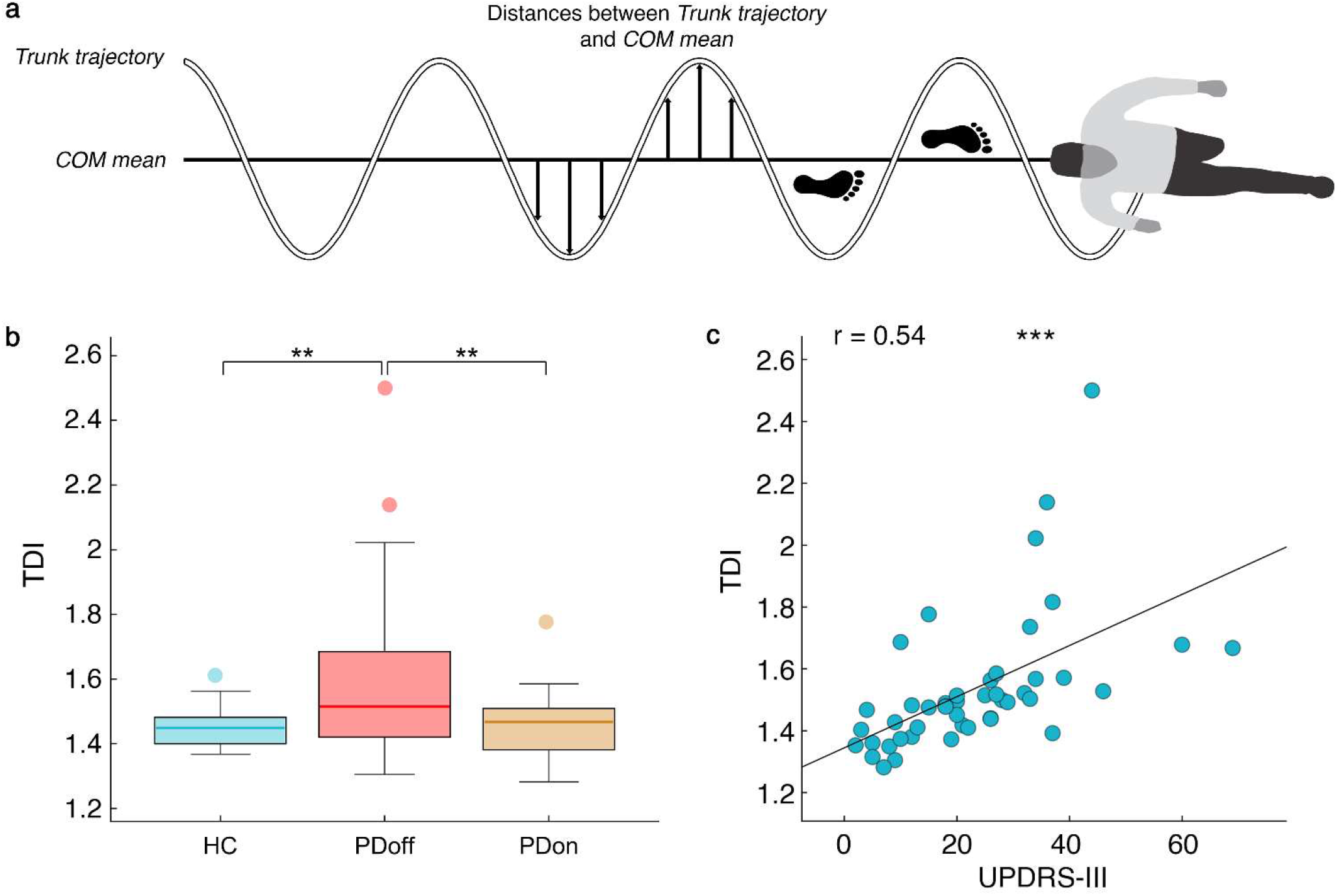
Trunk displacement index (TDI). **a.** The graphical representation of a step of the TDI calculation. The horizontal line is the mean position of the centre of mass on the x axis (COM mean). The waving line is the trajectory of the trunk. The arrows represent the distance between trunk trajectory and COM mean in each frame. **b.** The box plot (see Fig. 1 caption for explanation of box plot) of the TDI comparison among healthy controls (HC), individual with Parkinson’s disease before L-DOPA intake (PDoff) and individual with Parkinson’s disease after L-DOPA intake (PDon). **c.** The correlation between TDI and Unified Parkinson’s Disease Rating Scale Part III score (UPDRS-III), including PDoff and PDon. Significance *p* value: \**p*<0.05, \*\**p*<0.01, \*\*\**p*<0.001.

### 2.4 Statistical analysis

The statistical analysis was performed in MATLAB, (Mathworks®, version R2018b). In order to compare our data, we performed permutation tests, in which each subject label has been permuted 10,000 times. The comparison among the three groups was carried out through permutational multivariate analysis of variance (PERMANOVA) to obtain a null distribution. A Pearson’s correlation analysis was performed to determinate the association between TDI and the motor evaluations, regardless of the group.

All the reported *p*-values have been corrected for multiple comparisons with the false discovery rate (FDR)^31^. A significance level of p<0.05 has been considered.

## 3 RESULTS

### 3.1 Demographic, clinical, neuropsychological evaluation

In the present study, we evaluated 23 PD patients (in on and off state) and 23 healthy controls. As shown in Table 1, no statistically significant differences have been found for demographic, anthropometric and neuropsychological data except for the UPDRS-III in the PD, between off and on condition, with the PDoff group showing higher UPDRS-III values.

**Table 1.**
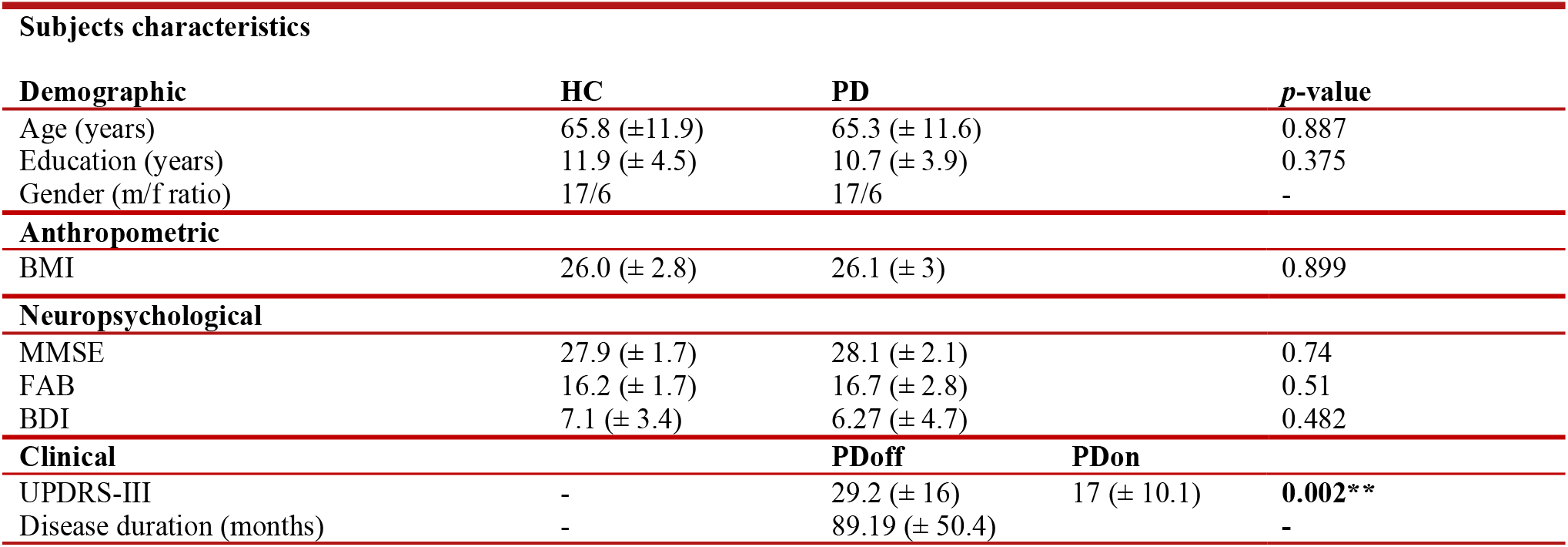
Comparison between healthy control group (HC) and individuals with Parkinson’s disease (PD), for demographic, anthropometric, neuropsychological parameters. Clinical variables have been compared within the PD group before (PDoff) and after (PDon) L-DOPA intake. Body mass index (BMI), mini mental state examination (MMSE), frontal assessment battery (FAB), Beck’s depression inventory (BDI), unified Parkinson’s disease rating scale part III (UPDRS-III). Value expressed as mean (± standard deviation). Significance *p* value: * *p*<0.05, \*\**p*<0.01, \*\*\**p*<0.001.

### 3.2 Motion evaluation (before and after L-DOPA intake)

In order to assess the sensitivity of the calculated parameters, a comparison among the three groups (HC, PDoff, PDon) has been performed.

Spatio-temporal comparison among the three groups showed statistically significant differences in variability parameters (Figure 3). In particular, compared to both PD groups, the HC group showed low CV values of stride time (HC vs PDoff, p=0.003; HC vs PDon, p=0.006), swing time (HC vs PDoff, p<0.001; HC vs PDon, p=0.039), DLS (HC vs PDoff, p<0.001; HC vs PDon, p=0.009) and stride length (HC vs PDoff, p=0.001; HC vs PDon, p=0.051). The remaining spatio-temporal parameters failed to show any statistically significant difference. None of the spatio-temporal parameters succeeded in distinguishing the PDoff group from the PDon group.

**Figure 3.**
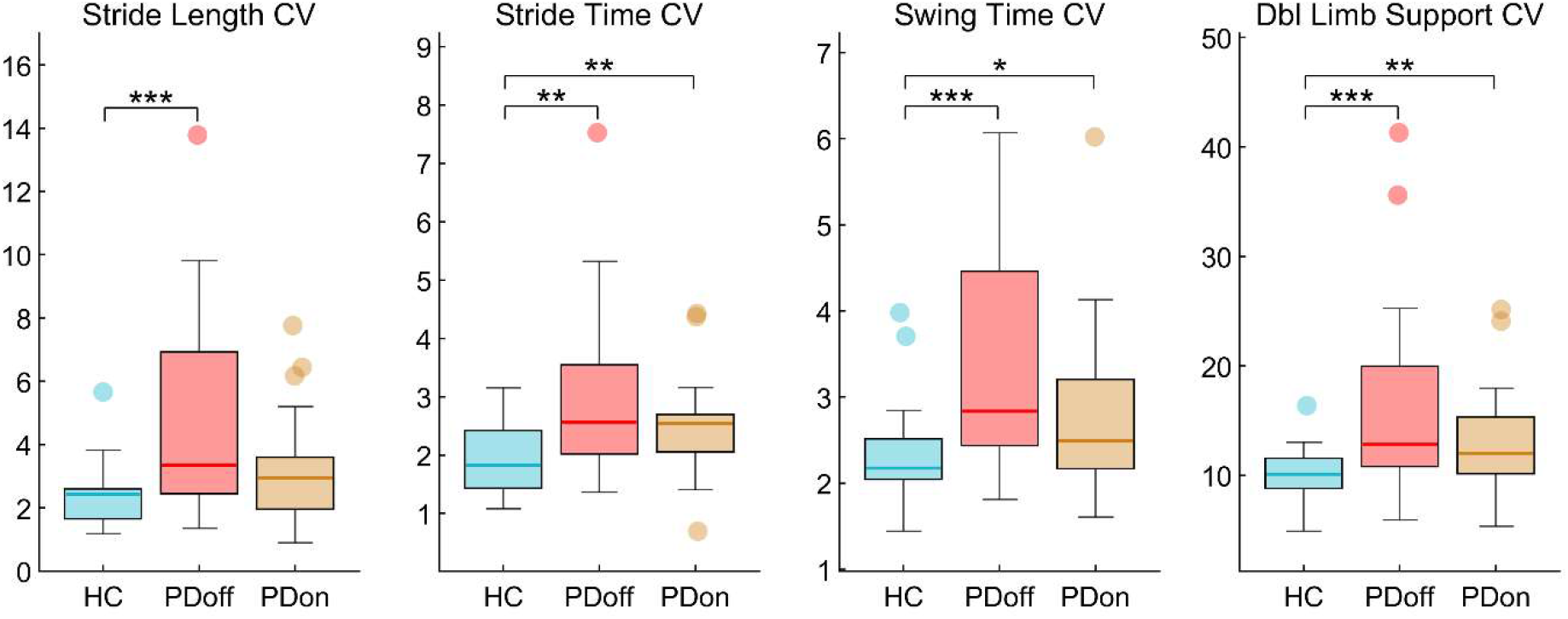
Spatio-temporal analysis of gait. Box plots (see Fig. 1 caption for explanation of box plot) of coefficients of variability (CV) of spatio-temporal parameters. Healthy controls (HC), individual with Parkinson’s disease before L-DOPA intake (PDoff), individual with Parkinson’s disease after L-DOPA intake (PDon). Significance *p* value: \**p*<0.05, \*\**p*<0.01, \*\*\**p*<0.001.

Kinematic analysis among the three groups (Figure 1) displayed statistically significant differences in the ankle ROM values. Specifically, the HC group showed high AΔ1 values compared to both PD groups (HC vs PDoff, p<0.001; HC vs PDon, p=0.014). Not even the kinematic parameters could recognise any difference between the off and on condition in PD group.

Concerning the statistically significant parameters, PDon values of CV and articular ROM have always resulted halfway between HC group and PDoff group.

Finally, in the TDI analysis (Figure 2b) the PDoff group displayed high TDI values compared to both HC (p=0.005) and PDon (p=0.004) groups. The TDI successfully distinguished the PD group before and after L-DOPA subclinical treatment.

### 3.3 TDI correlation analysis

In order assess the biomechanical and clinical significance of the trunk displacement index, we carried out a correlation analysis examining the TDI relationship with gait parameters (spatio-temporal, CV and kinematic) and clinical assessment (UPDRS-III). Figure 4 shows the correlation plots of TDI with spatio-temporal parameters. We found positive correlations between TDI and stability parameters such as stride width (r=0.28, p=0.03), stride length CV (r=0.74, p<0.001), stride time CV (r=0.61, p<0.001), swing time CV (r=0.53, p<0.001), cycle time CV (r=0.64, p<0.001), strides/minute CV (r=0.63, p<0.001) and DLS CV (r=0.53, p<0.001). Furthermore, we found negative correlations between TDI and velocity parameters such as speed (r=−0.68, p<0.001) and stride length (r=−0.73, p<0.001).

**Figure 4.**
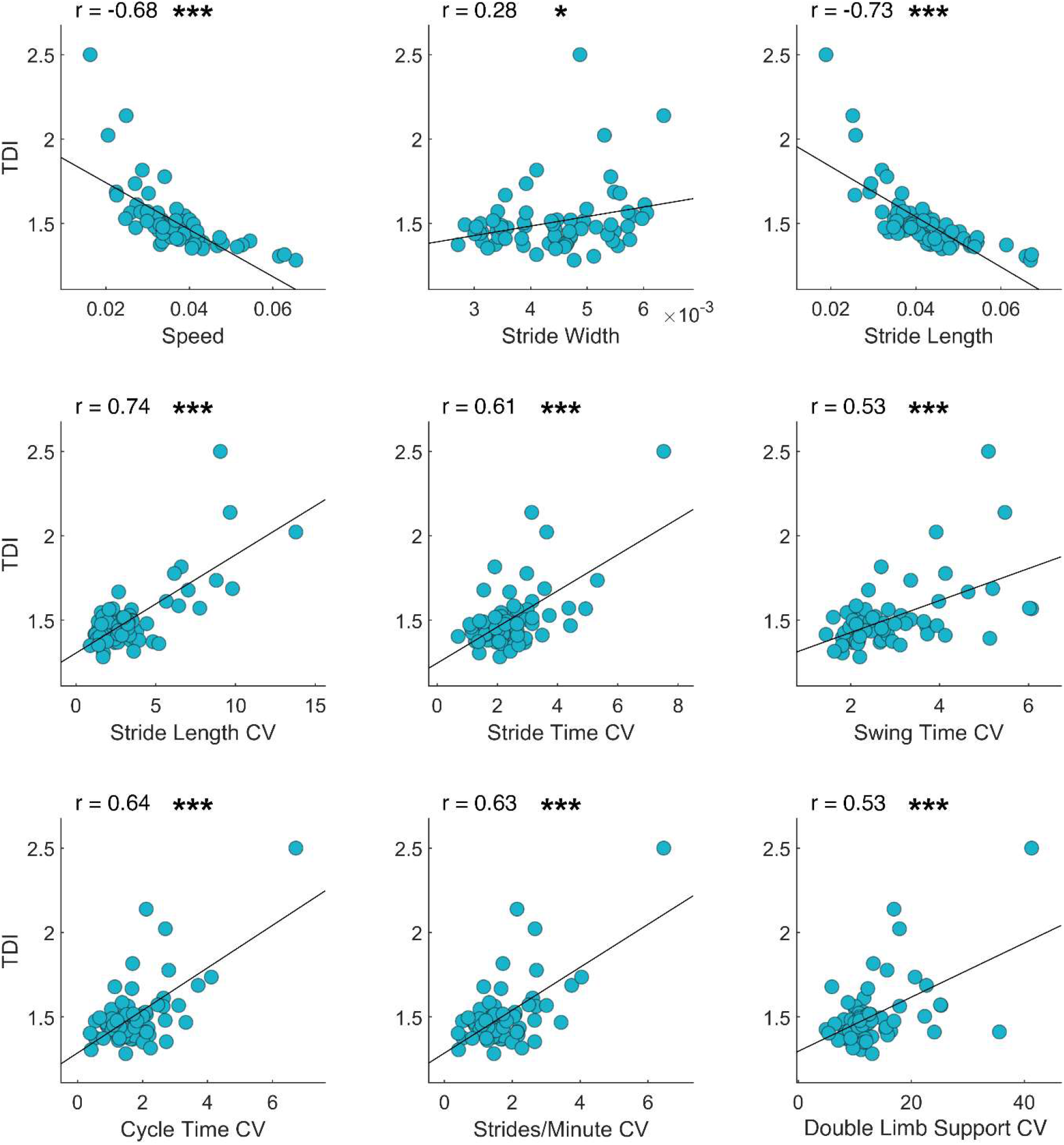
TDI and spatio-temporal gait parameters. Pearson coefficient correlation between trunk displacement index (TDI) and spatio-temporal gait parameters. The correlation includes healthy controls, individuals with Parkinson’s disease before L-DOPA intake, individuals with Parkinson’s disease after L-DOPA intake. Speed (m/s), Stride width (m), Stride length (m), Coefficient of variability (CV). Significance *p* value: \**p*<0.05, \*\**p*<0.01, \*\*\**p*<0.001.

TDI and kinematic parameters always showed negative correlations (Figure 5): AΔ1 (r = −0.45, p < 0.001), AΔ3 (r=−0.66, p<0.001), AΔ4 (r=−0.71, p<0.001), KΔ1 (r=−0.59, p<0.001), KΔ2 (r=−0.4, p=0.001), KΔ3 (r=−0.57, p<0.001), KΔ4 (r=−0.6, p<0.001), TΔ2 (r=−0.61, p<0.001), TΔ3 (r=−0.61, p<0.001).

**Figure 5.**
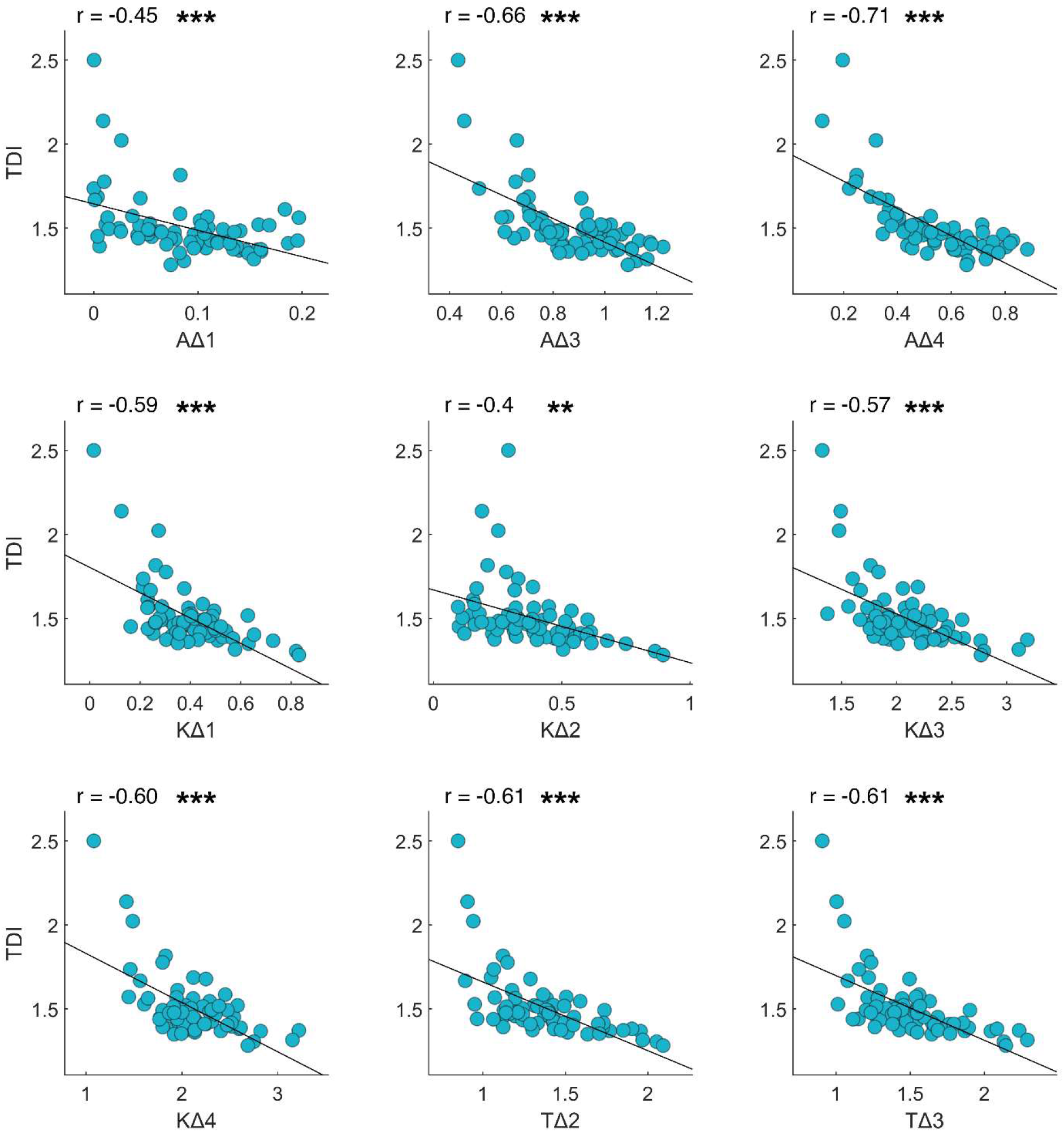
TDI and kinematic gait parameters. Pearson coefficient correlation between trunk displacement index (TDI) and kinematic gait parameters. The Δ values are calculated as the difference between two consecutive peaks (minimum or maximum joint excursion degree). The correlation includes healthy controls, individuals with Parkinson’s disease before L-DOPA intake, individuals with Parkinson’s disease after L-DOPA intake. Ankle Δ (AΔ−), Knee Δ (KΔ−), Thigh Δ (TΔ−). Significance *p* value: \**p*<0.05, \*\**p*<0.01, \*\*\**p*<0.001.

Finally, Figure 2c shows the positive correlation between TDI and clinical motor condition assessed trough the UPDRS-III score (r = 0.538, p < 0.001).

## 4 DISCUSSION

In this study we extracted from 3D-GA data, a biomechanical index, named TDI, capable of synthetically conveying the complex motor impairment of PD patients. The TDI was obtained as the ratio between trunk and COM displacement and was correlated to both the typical gait parameters and UPDRS-III score. In order to assess its sensitivity, the ability to discriminate the motor effects of a subclinical dose of L-DOPA intake was evaluated.

### 4.1 Motion assessment and sensitivity evaluation

#### 4.1.1 Spatio-temporal assessment

The 3D-GA of spatio-temporal parameters revealed that both PD groups showed higher CV in stride time, swing time and DLS, compared to the HC group. Moreover, stride length CV was statistically different only when comparing the PDoff and HC groups, although when comparing the PDon and HC groups a strong statistical tendency was evident (p = 0.051). The higher variability of gait parameters may reflect the well-known loss of harmonic motor control that characterizes PD^32^. Our findings are in agreement with studies reporting higher variability in PD patients, compared to healthy age-matched controls^33,34^.

Despite a slight decrease, CVs of PDon group did not show any statistical difference from PDoff group, remaining statistically higher than HC group. These results may be interpreted both as a lack of sensitivity of CV parameters in recognising subclinical L-DOPA dose effects and as a persistence of the motor impairment after subclinical medication. Our results agree with studies reporting a high variability of gait in PD patients, even after L-DOPA intake^34–36^.

Beyond the CV parameters, the lack of significant differences in the remaining spatio-temporal parameters among the three groups, were similarly reported in studies investigating early PD patients^37,38^. However, other studies on analogous populations were able to find significant differences in the spatio-temporal parameters, with the PD patients showing reduced velocity characteristics of gait^39,40^. The difference with our findings, may be due to a milder motor impairment of our early PD patients.

#### 4.1.2 Kinematic assessment

With regard to the kinematics parameters, both PD groups, compared to the HC group, showed a lower ankle ROM at the beginning of the gait cycle (initial contact and load response), consistent with a significant reduction of the stride length. These results are in line with previous evidence about articular kinematics in PD^41^. However, some authors reported a reduction of ROMs also in knee and thigh^14,42,43^. It is noteworthy that following L-DOPA intake, ankle ROM, despite a slight increase, has remained significantly lower when compared to HC group. Similar results have been reported by Wu et al, showing that after pharmacological treatment, even if PDon ankle ROM increased, differences from HC group were still present, especially during the heel strike^40^.

The comparison between the on and off condition, for both spatio-temporal and kinematic parameters, failed to show any significant modification induced by subclinical L-DOPA doses.

#### 4.1.3 TDI assessment

Unlike the spatio-temporal and kinematic gait parameters, the TDI was able to distinguish the PDoff group not only from healthy controls but also from the PDon group. These data suggest the high sensitivity of the TDI, being a measure able to detect the slight effects of a subclinical dose of L-DOPA on the overall motion features.

Trunk displacement in PD is generally analysed with methodological approaches different from 3D-GA. Adkin et al. employed angular-velocity transducers to measure trunk sway in PD patients (off and on conditions), and healthy age-matched controls during gait. The results could not show any medication effect, but highlighted the presence of an impaired trunk mobility^44^. Mancini et al., evaluating trunk sway through jerk (the first time derivative of acceleration) in naïve PD patients, showed higher medio-lateral sway in PD group compared to healthy controls^45^. Horak et al., conducted an analysis of the trunk domain (comprehending ROM, peak velocity and acceleration) evaluating PD patients (off and on conditions) and healthy controls. The trunk domain in PD patients, despite a significant improvement after L-DOPA intake, remained statistically different when compared to healthy controls^46^. Cole et al., using electromyography, analysed trunk stability and displacement in PD subjects with a history of falls. The study reported a greater trunk muscle activation in PD patients, causing impaired trunk control^47^. Despite methodological differences, all these studies agree about the existence of a trunk impairment in PD. Our analysis, besides being consistent with this observation, showed that the TDI resulted sensitive enough to distinguish patients in off and on conditions. Furthermore, the dimensionless quality of the TDI offers the advantage of a measure which does not depend on time and makes acquisition of different duration easily comparable. It is equally important that such an index takes into account not only the trunk, but also the centre of mass, a biomechanical element deeply involved in balance dynamics^48^ and suprasegmental control of stability^18^. Moreover, a local reference point such as the COM mean, rather than a global one, allows a displacement evaluation not conditioned by external variables.

### 4.2 The biomechanical and clinical significance of the TDI

#### 4.2.1 TDI and spatio-temporal parameters

Our analysis showed negative correlations with velocity parameters such as speed and stride length, implying that when TDI decreases (HC and PDon groups have lower TDI) speed and stride length increase. The gait pattern adopted by PDoff patients, characterized by a lower walking speed and a shorter stride length, both associated with a higher TDI, may be interpreted as a precautionary strategy to reduce biomechanical distresses and risk of falls^49^. Walking speed has been correlated with fall risk several times^50–53^. Furthermore, according to Bayle et al., the walking speed is influenced by the stride length, and this parameter displays a negative correlation with the time from the clinical onset, making it a clinical marker for PD^54^. This result is in accordance with our finding of a negative correlation of TDI with stride length, where low displacement values correlated with higher stride length.

Furthermore, TDI correlates positively with stride width and with the CV of many spatio-temporal parameters, a variable known to be predictive of fall^49,55–57^. Observing the positive correlations of TDI with the variability of many gait parameters, we supposed that the TDI may represent an aspect of the suprasegmental control mechanism of stability. However further analysis with specific design are needed in order to demonstrate this characteristic.

#### 4.2.2 TDI and kinematic parameters

From the correlation analysis between articular kinematic parameters and TDI it emerged that 9 (out of 12) ROMs (AΔ1, AΔ3, AΔ4, KΔ1, KΔ2, KΔ3, KΔ4, TΔ2, TΔ3) showed a negative correlation with TDI. According to this observation, the index resulted able to represent the articular mobility of lower limbs, in which a lower TDI is related to higher mobility and vice versa. Studies generally agree about the fact that the joint ROMs are larger in the healthy population than in PD patients^40,41^. In particular, Morris et al., in a study on kinematics of gait, showed that PD patients in the off condition displayed the lowest ROM degrees in all three main articulation of lower limbs (ankle, knee, thigh); the same patients, in the on condition, showed a slight increase of all three ROMs, but healthy control group remained the group with the highest ROMs in all the joint considered^14^.

#### 4.2.3 TDI and UPDRS-III

Finally, another very intriguing result is represented by the correlation that TDI shows with the motor UPDRS-III. The highly statistically significant correlation with UPDRS-III makes the TDI an efficient, global marker of the motor condition in the PD patient. Indeed, the positive correlation shows that when the trunk displacement index grows (PDoff group has higher TDI), the UPDRS-III score grows accordingly (worse motor condition). This finding suggests that the TDI, an objective measure, may be able to compensate for the limitations of the UPDRS. Moreover, it could be used in accurate clinical evaluations of the overall motor condition of PD patients, even to assess the effects of therapeutic strategies.

### 4.3 Conclusion

In this first study on the TDI, the high number of significant correlations and the coherence of the coefficients with the gait parameters reflects the intrinsic characteristics of the index, providing it with a clear biomechanical significance. In addition, the correlation with the UPDRS-III highlights its clinical relevance in PD motor evaluation. Hence, the TDI may be able to offer both a synthetic expression of the motor condition and a representation the balance control of the PD patients. Moreover, the TDI displays a high sensitivity. In fact, despite the subclinical L-DOPA dose and the short wearing off time, which prevents the elimination of the long-duration effects of L-DOPA^58^, the TDI could still capture significant difference between the off and on conditions.

Summarising, the TDI reflects the overall motor condition of patients with PD in a very effective and sensitive way. To this respect, the TDI may offers a new and improved tool to analyse gait control following pharmacological or rehabilitation protocols.

